# Whole-body homogenates restore disrupted microbiota composition in a model insect better than feces or no restoration treatment

**DOI:** 10.1101/2025.11.17.688872

**Authors:** Ricardo Romo, Erin Ricks, Reed Ogden, Paige E. Bonnette, Shelby C. Olson, Krista Pearman, Gerald B. Call, John M. Chaston

## Abstract

Antibiotic treatment can disrupt gut microbiota and pose challenges and opportunities for the establishment or restoration of healthy microbial communities. Using the fruit fly, *Drosophila melanogaster*, as an experimental model, we evaluated the impact of two types of microbial transplants—fly feces and whole-body fly homogenates—on host microbiota composition, following, or independent of, tetracycline-induced community disruption. Using 16S rRNA sequencing, we compared community beta diversity between treatments. We show that antibiotic treatment significantly altered microbiota composition and community structure relative to untreated controls. Flies inoculated with whole body homogenates of age-matched, antibiotic-free flies had a more similar microbial community composition to the untreated communities than flies exposed to fly feces or to flies that received no restoration treatment. We also found that the presence of *Wolbachia* was associated with variation in microbiota composition and specific locomotor functions. These findings show that whole-body homogenates are a superior method for microbiota restoration in *Drosophila melanogaster* and contribute to a growing body of research on microbial community restoration following disturbance.

**IMPORTANCE:** Gut microbes play a critical role in animal biology, influencing digestion, immunity, development, and behavior. Disruptions to the gut microbiota—whether from antibiotics, disease, or other interventions—can have lasting effects, and restoring these communities remains an important challenge across biological and biomedical research. Model organisms like fruit flies (*Drosophila melanogaster*) provide a powerful system for testing microbial restoration methods because their gut communities are relatively simple and easy to manipulate. In this study, we compared various strategies for repopulating the microbiota of flies following antibiotic treatment. We found that flies fed whole-body homogenates of untreated flies more closely resembled the microbiota of untreated flies than flies exposed to fly feces or to flies that received no restoration treatment.

These findings contribute to broader efforts to understand and develop methods for microbiota recovery following disturbance and suggest applications across animal systems.

## INTRODUCTION

Animal-associated microbial communities help shape host health by contributing to digestion, immune development, metabolism, and behavior (1, 2). Disruptions to the microbiota caused by antibiotics, disease, or environmental change can lead to physiological consequences, including inflammatory and neurodevelopmental conditions (3, 4). As a result, microbiota restoration is a growing area of interest in both clinical and experimental contexts.

The gut microbiota of the fruit fly, *Drosophila melanogaster*, is of relatively low diversity and abundance, making it a useful system for studying host–microbe interactions (5, 6).Microbial colonization of adult flies is maintained by continuous exposure to environmental microbes through diet and frass (7, 8). The fly microbiota is generally dominated by Lactic Acid Bacteria and Acetic Acid Bacteria, especially members of *Lactobacillus* and *Acetobacter*, which are known to impact host development and metabolism (9, 10). Flies can be rendered germ-free by bleaching embryos to remove the chorion, enabling researchers to raise them under gnotobiotic conditions and introduce defined microbial communities in a controlled manner (11). This approach allows precise testing of how individual bacterial species or synthetic communities influence host traits (9). Therefore, experimental control of the microbiota is relatively straightforward in *Drosophila*.

Restoration of disrupted microbial communities is a relatively understudied area in *Drosophila*, possibly because gnotobiotic approaches are more suitable for most research questions being addressed in fruit flies. Antibiotics have sometimes been administered to fruit flies to eliminate, control, or, more commonly, disrupt, their microbiota (12-19). Among these studies, various metrics have been used to gauge the effectiveness of antibiotic treatment, but most analyses suggest antibiotic treatment likely alters but does not completely eliminate associated microorganisms from adult flies unless it is combined with other treatments, such as sterilization or removal of the embryonic chorion. These examples highlight that antibiotic treatment has been widely, but inconsistently, used as a substitute for axenic rearing, underscoring both its experimental utility and limitations.

One area where antibiotic treatment is necessary in flies, is to eliminate the reproductive endosymbiont, *Wolbachia. Wolbachia* are maternally transmitted, intracellular bacteria that infect a wide range of arthropods, including many *Drosophila* species (20). *Wolbachia* can manipulate host reproduction through mechanisms such as cytoplasmic incompatibility, male killing, feminization, and parthenogenesis, thereby spreading rapidly in host populations (20-23). In addition to reproductive effects, *Wolbachia* can alter host immunity, metabolism, and even viral resistance (24, 25). When flies are reared for 2–3 generations on diets containing tetracycline, *Wolbachia* are readily eliminated from subsequent generations (25-30). Although effective in eliminating *Wolbachia*, the effects of tetracycline exposure can also perturb the gut microbiota (19). However, the influence of multi-generational exposure to tetracycline for the elimination of *Wolbachia*, and whether this influences the restoration of a disturbed microbial community, has not been investigated previously.

Several studies have tried to control for variation in fruit fly microbiota composition using restoration-style approaches. Restoration is often necessary because flies acquire their microbes from their environment leading to inconsistencies in microbial community composition following disruption (8, 31). Researchers have explored non-gnotobiotic restoration methods following antibiotic disruption, including exposing flies to donor fecal material (16, 32) or whole-body or gut homogenates from untreated donors (33). Here, we investigate microbiota restoration in antibiotic-treated flies, testing whether whole-body homogenate feeding or feces transfer more effectively restores microbial composition. Using 16S rRNA sequencing and beta diversity analysis, we identify which approach best approximates the microbiota of untreated flies. We also examine how *Wolbachia* status influences restoration outcomes and, when comparing fly populations with comparable microbiota composition, specific metrics of fly locomotor activity. These findings contribute to broader efforts to understand microbiota recovery dynamics and offer guidance for experimental reproducibility in host–microbiota studies.

## METHODS

### Fly rearing methods and sample collection

All flies were derived from a *w*^*1118*^ laboratory stock of *D. melanogaster* that harbored *Wolbachia*. Standard rearing conditions were at 25°C on a 12-hour light:dark cycle at ambient humidity on a cornmeal diet (411.8 g cornmeal (Agricor 104CMYLWCR), 247 mL light corn syrup (Karo), 247.1 mL molasses (Grandma’s Unsulphured), 82.4 g yeast (Red Star Active Dry), 41.2 g soy flour (Genesee Scientific 62.115), 41.2 g agar (Genesee Scientific 66-104), 30.9 mL propionic acid, 70 mL 10% methylparaben in 95% ethanol). Flies were divided into two main categories that were maintained with the same generation times as our main stock, which we call the **Untreated** population (**FIG 1**). These categories were *Wolbachia-*positive (W+) *and Wolbachia-*negative (W-). Four independent W-populations were established by placing Untreated W+ adult flies on standard diet that was supplemented with 30 μg/ml tetracycline. After these Parental (P) generation flies laid embryos for several days, they were removed from the vials. Eclosed offspring were transferred to fresh tetracycline-supplemented diet as described for two more generations for a total of three sequential generations exposed to tetracycline. Eclosing F_3_ flies were collected to vials of standard diet lacking tetracycline and, after laying embryos for several days, were tested for the presence of *Wolbachia* by PCR. DNA was extracted from representative flies using the Zymo Fecal/Soil DNA kit. The presence of *Wolbachia* was detected by PCR using wsp691-R (5’-AAAAATTAAACGCTACTCCA-3’) and wsp81-F (5’-TGGTCCAATAAGTGATGAAGAAAC-3’) primers as described previously (34), We used NEB Taq polymerase (M0267S) with 20 μl reaction mixtures, including 10X Thermopol Buffer, 25 mM MgCl_2_, 10mM each dNTP, 20mM each primer, and 1 μl template DNA. Thermal cycler settings were at 94°C for 3 min, 35 cycles of 94°C for 1 min, 55°C for 1 min, and 72°C for 1 min, with a final 72°C extension at 3 min. The lines were considered *Wolbachia*-free if there was no ∼ 500 bp band on the gel. Four healthy independent W-lines were selected for future analysis (1-4), and reared on tetracycline-free diet for two generations. When these F_5_ flies were five days post eclosion, they were transferred to standard diet to lay F_6_ embryos in vials prepared in advance with microbiota restoration treatments. For the **Feces** treatment, standard diet was pre-seeded for three days with 40 male W+ flies (males deposit frass but do not lay embryos). For **Homogenate** restoration, vials were inoculated with 60 μl of a whole-body male fly homogenate and air dried in a biosafety cabinet. The homogenate was prepared from ten F_5_ untreated male flies in 250 μl of PBS by beating CO_2_ anesthetized flies with lysis beads in a Benchmark D1030 BeadBug Microtube Homogenizer at 400 (Speed x 10) for 45 s. For the **No Restoration** condition, standard diet vials received no microbial pretreatment. At the same time, four replicate populations of adult W+ flies were subjected to each of the same treatments (Populations 5-8) so that all eight populations (four W- and four W+) received each of the three restoration conditions (No Restoration, Feces, Homogenate) in separate, replicated vials. At the time the F_6_ populations were established we transferred ten pools of five F_5_ Untreated male flies and five pools of five F_5_ Untreated female flies to -80°C following light CO_2_ anesthesia, to represent the original target state of the restoration.

**Figure 1.**
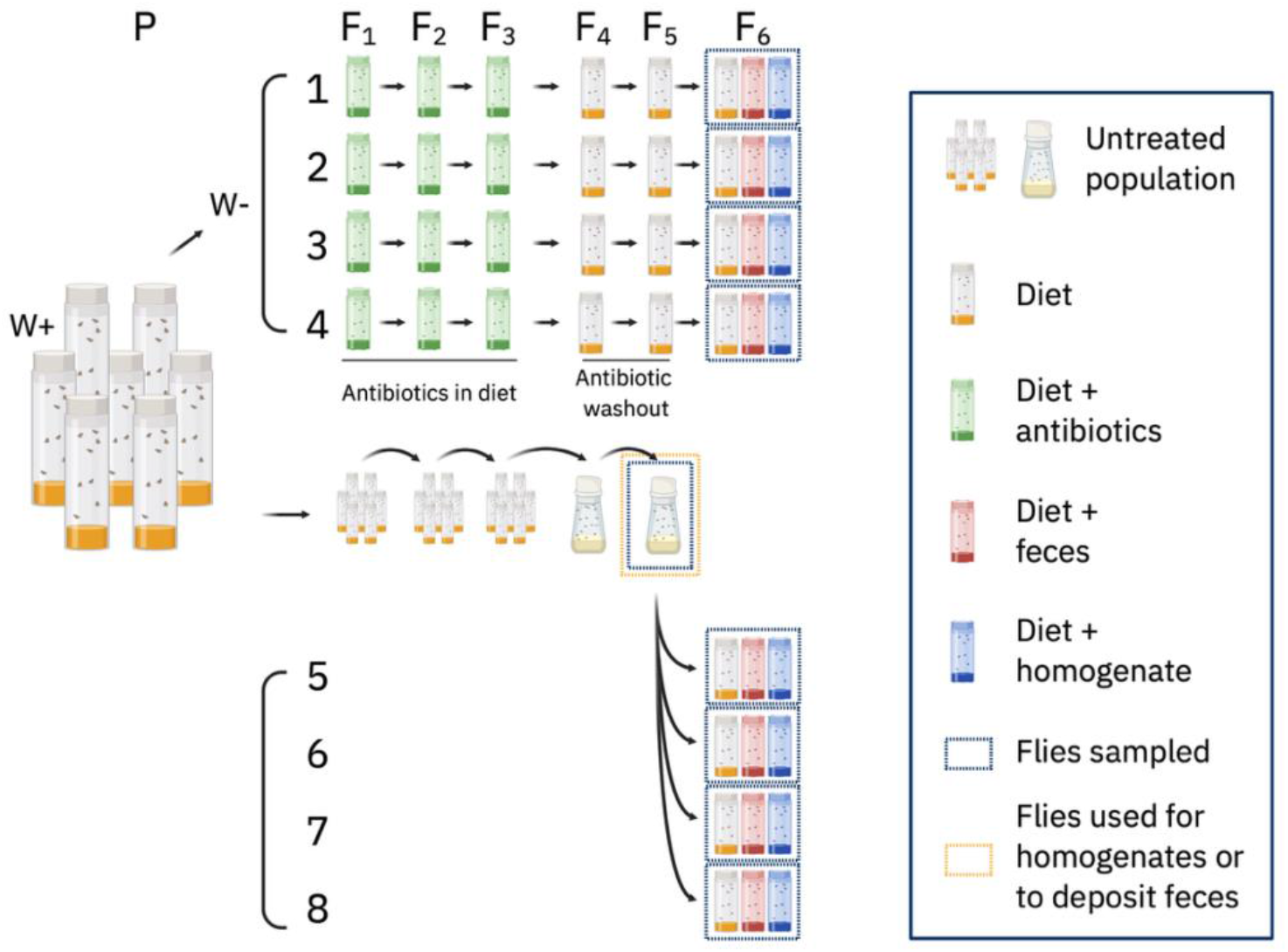
Schematic to prepare 8 distinct fly populations. From the Untreated W+ population, four W-populations (1-4) were derived by rearing on antibiotics for three generations and then on standard diet for two generations. From F_5_ Untreated flies, four replicate W+ populations (5-8) were established that were age-, generation-, and density-matched to Populations 1-4. Populations 1-8 were reared on standard diet alone, or preseeded with adult fly homogenate or adult fly feces. The homogenate and feces were derived from the same F_5_ population as was used to establish W+ populations 5-8. Both bottles and multiple vials are used to show Untreated flies. This best represents how they were maintained in culture and visually distinguishes Untreated flies in bottles from antibiotic washout and No Treatment flies in vials.

When the F_6_ flies were seven days post-eclosion, we collected ten pools of five flies (5 pools each of male and female) for every condition (eight lineages * three conditions * ten replicates = 240 samples). Each sample was stored at -80°C in preparation for DNA sequencing.

### Sample preparation and DNA sequencing

DNA was extracted from each sample using a Zymo Fecal/Soil Microbe 96 Kit and following manufacturer instructions, including adding ß-mercaptoethanol to the ZR BashingBead™ Lysis Buffer. We included reagent-only control blanks, and the ZymoBIOMICS Microbial Community DNA Standard as a positive control. DNA sequencing libraries were prepared as described previously (35). Briefly, we used Accuprime Pfx DNA Polymerase, and PCR cycling conditions were: 95 °C for 120s; 30 cycles of 95 °C for 20s, 55 °C for 15s, and 72 °C for 300s; and a final extension at 72 °C for 600s. We confirmed via gel electrophoresis that there was no amplification of the negative controls. Then, samples were normalized using the Just-a-Plate™ 96 PCR Normalization and Purification Kit (Charm Biotech) and pooled in batches of 96 samples. Each pool was concentrated to 10 ml using the DNA Clean & Concentrator®-25 kit (Zymo Research). A BluePippin size selection step was performed to remove fragments outside the 250–450 bp range. The samples were then sequenced on a partial 500-cycle Illumina MiSeq lane at the Arizona State University Genomics Core Facility, which yielded a total of 11,999,490 reads across 261 samples (median 27,284, min 3, max 412392 reads per sample). At each nucleotide position, 1^st^ quartile forward and reverse read quality scores were above 30.

### Bioinformatics and Data Analysis

Microbiota data were processed using QIIME2 version 2024.10 (36) and DADA2 (37) in a reproducible Conda environment. After confirming *Wolbachia* reads were absent in the antibiotic-treated samples, we filtered all reads that could be assigned to *Wolbachia, Chloroplast, Mitochondria, Rickettsia*, and *Archaea*, and rarefied OTU tables to a depth of 500 reads per sample. This left a total of 127,500 reads across 255 samples comprising 171 amplicon sequence variants (ASVs). We also confirmed that rarefied read counts approached saturating sampling based on the rarefaction curve (**FIG S1**). Phylogenetic trees were generated using MAFFT alignment (38) and FastTree inference (39) and were incorporated into phylogeny-aware distance metrics. Beta diversity was assessed using Bray-Curtis dissimilarity, weighted UniFrac, and unweighted UniFrac distances (40, 41). Principal Coordinates Analysis (PCoA) was performed on each distance matrix. Subsets of the rarefied feature table were created to compare treatment groups. PERMANOVA was performed with 999 permutations (42). Taxonomic profiles were visualized as ASV-level bar plots using ggplot2 (43). Differential abundance testing was conducted at the ASV level using standard ANCOM (44). Statistical analyses and visualizations were performed in R or QIIME2.

### Climbing Assay

The climbing assay was performed as previously described (45). Briefly, 10-day-old flies were individually loaded into climbing vials using a fly aspirator. Loaded climbing vials were then placed vertically into a multibeam monitor (MBM, TriKinetics) and fly position was monitored for 20 minutes. Data was analyzed in Excel and GraphPad Prism 10. Data represent the mean ± standard error of the mean and unpaired t-tests were performed.

## RESULTS

### Whole-body adult homogenates produce a native-like microbiota in conventionally-reared flies

We first examined how different microbiota treatment strategies influenced microbiota composition in flies with an established microbiota. Consistent with previous demonstrations in laboratory flies, the microbiota of the flies we examined was of low diversity; in this case, the fly microbiota was dominated by just four ASVs, one each assigned as *Acetobacter* and *Commensalibacter*, and two assigned as *Lactiplantibacillus plantarum* (**FIG 2A-M**). Microbiota composition varied significantly between flies that received one of three treatments: 1) Feces or 2) a whole-body Homogenate of age- and generation matched Untreated flies in the immediately preceding generation; or 3) No Restoration treatment (PERMANOVA F _3, 116_ = 17.50, R^2^ = 0.27, p = 0.001). Bray-Curtis and Unifrac distance analysis confirmed that homogenate treated flies had smaller differences relative to and were more frequently similar to Untreated flies in the immediately preceding generation (**FIG 2N, FIG S2A-C, TABLE 1**). The greater similarity of Homogenate-treated flies to Untreated flies, relative to the No Restoration treatment, is consistent with inconstant variation in microbial community composition between generations and a microbiota composition that is driven by dietary replenishment. (8, 31). Community level variation was likely driven at least in part by differences in the abundances of a *Commensalibacter* ASV, the only ASV that varied significantly between treatments as reported by ANCOM. Taken together, these findings support previous evidence that the microbiota of *Drosophila* is inconstant and show that microbiota manipulations can maintain microbiota consistency across generations. We conclude that inoculating flies with a whole-body homogenate provides better generational maintenance of the microbiota than exposure to feces or cooked (but not sterilized) diets.

**Table 1.** PERMANOVA results comparing microbial beta diversity across Untreated, W^+^, and W-samples under different microbiome restoration treatments.

|  | Bray Curtis |  |  |  |  | Weighted Unifrac |  |  |  | Unweighted Unifrac |  |  |  |
| --- | --- | --- | --- | --- | --- | --- | --- | --- | --- | --- | --- | --- | --- |
|  | df<sup>a</sup> | SS<sup>b</sup> | R<sup>2</sup> | F | p<sup>c</sup> | SS | R<sup>2</sup> | F | p | SS | R<sup>2</sup> | F | p |
| W+ treatments and Untreated |  |  |  |  |  |  |  |  |  |  |  |  |  |
| Treatment | 3 | 0.98 | 0.19 | 18.2 | 0.001 | 0.14 | 0.4 | 29.9 | 0.001 | 0.045 | 0.05 | 2.3 | 0.009 |
| Sex | 1 | 0.15 | 0.028 | 5.5 | 0.006 | 0.0025 | 0.007 | 1.6 | 0.18 | 0.062 | 0.07 | 9.5 | 0.001 |
| Population (1-4) | 3 | 0.55 | 0.11 | 6.8 | 0.001 | 0.027 | 0.078 | 5.9 | 0.001 | 0.022 | 0.025 | 1.1 | 0.3 |
| Residual | 116 | 3.1 | 0.60 | - | - | 0.18 | 0.52 | - | - | 0.76 | 0.85 | - | - |
| Total | 123 | 5.2 | 1 | - | - | 0.35 | 1 | - | - | 0.89 | 1 | - | - |
| W- treatments and Untreated |  |  |  |  |  |  |  |  |  |  |  |  |  |
| Treatment | 3 | 2.2 | 0.67 | 87.5 | 0.001 | 0.17 | 0.68 | 96.0 | 0.001 | 0.28 | 0.094 | 4.2 | 0.001 |
| Sex | 1 | 0.025 | 0.0076 | 3.0 | 0.072 | 0.0022 | 0.009 | 3.8 | 0.047 | 0.13 | 0.042 | 5.6 | 0.001 |
| Population (5-8) | 3 | 0.13 | 0.039 | 5.1 | 0.003 | 0.0093 | 0.037 | 5.3 | 0.001 | 0.06 | 0.02 | 0.9 | 0.61 |
| Residual | 113 | 0.95 | 0.29 | - | - | 0.067 | 0.27 | - | - | 2.5 | 0.84 | - | - |
| Total | 120 | 3.3 | 1 | - | - | 0.25 | 1 | - | - | 3 | 1 | - | - |
| W+ vs W- No Restoration (Populations 1-8) |  |  |  |  |  |  |  |  |  |  |  |  |  |
| Wolbachia | 1 | 1.638 | 0.480 | 63.5 | 0.029 | 0.135 | 0.520 | 72.7 | 0.250 | 0.194 | 0.155 | 13.2 | 0.001 |
| Sex | 1 | 0.084 | 0.025 | 3.2 | 0.036 | 0.002 | 0.008 | 1.2 | 0.288 | 0.076 | 0.061 | 5.2 | 0.001 |
| Wolbachia/Population (1-8) | 6 | 0.169 | 0.050 | 1.1 | 0.029 | 0.013 | 0.049 | 1.2 | 0.169 | 0.112 | 0.090 | 1.3 | 0.024 |
| Residual | 59 | 1.521 | 0.446 | - | - | 0.110 | 0.422 | - | - | 0.867 | 0.695 | - | - |
| Total | 67 | 3.412 | 1 | - | - | 0.260 | 1 | - | - | 1.249 | 1 | - | - |
| W+ vs W- Feces (Populations 1-8) |  |  |  |  |  |  |  |  |  |  |  |  |  |
| Wolbachia | 1 | 0.594 | 0.197 | 41.0 | 0.022 | 0.045 | 0.291 | 76.4 | 0.003 | 0.034 | 0.052 | 4.4 | 0.001 |
| Sex | 1 | 0.0509 | 0.020 | 4.1 | 0.041 | 0.007 | 0.043 | 11.2 | 0.003 | 0.052 | 0.080 | 6.9 | 0.001 |
| Wolbachia/Population (1-8) | 6 | 1.429 | 0.475 | 16.4 | 0.022 | 0.065 | 0.422 | 18.5 | 0.005 | 0.078 | 0.120 | 1.7 | 0.001 |
| Residual | 64 | 0.928 | 0.308 | - | - | 0.038 | 0.244 | - | - | 0.488 | 0.748 | - | - |
| Total | 72 | 3.010 | 1 | - | - | 0.154 | 1 | - | - | 0.652 | 1 | - | - |
| W+ vs W- Homogenate (Populations 1-8) |  |  |  |  |  |  |  |  |  |  |  |  |  |
| Wolbachia | 1 | 0.031 | 0.059 | 5.6 | 0.001 | 0.003 | 0.066 | 6.4 | 0.008 | 0.059 | 0.040 | 3.1 | 0.011 |
| Sex | 1 | 0.039 | 0.072 | 6.9 | 0.005 | 0.003 | 0.063 | 6.1 | 0.010 | 0.049 | 0.033 | 2.5 | 0.010 |
| Wolbachia/Population (1-8) | 6 | 0.113 | 0.211 | 3.4 | 0.001 | 0.009 | 0.220 | 3.5 | 0.007 | 0.157 | 0.106 | 1.4 | 0.094 |
| Residual | 63 | 0.351 | 0.658 | - | - | 0.027 | 0.652 | - | - | 1.215 | 0.821 | - | - |
| Total | 71 | 0.533 | 1 | - | - | 0.041 | 1 | - | - | 1.480 | 1 | - | - |

<sup>a</sup> degrees of freedom
<sup>b</sup> sum of squares
<sup>c</sup> p-value

**Figure 2.**
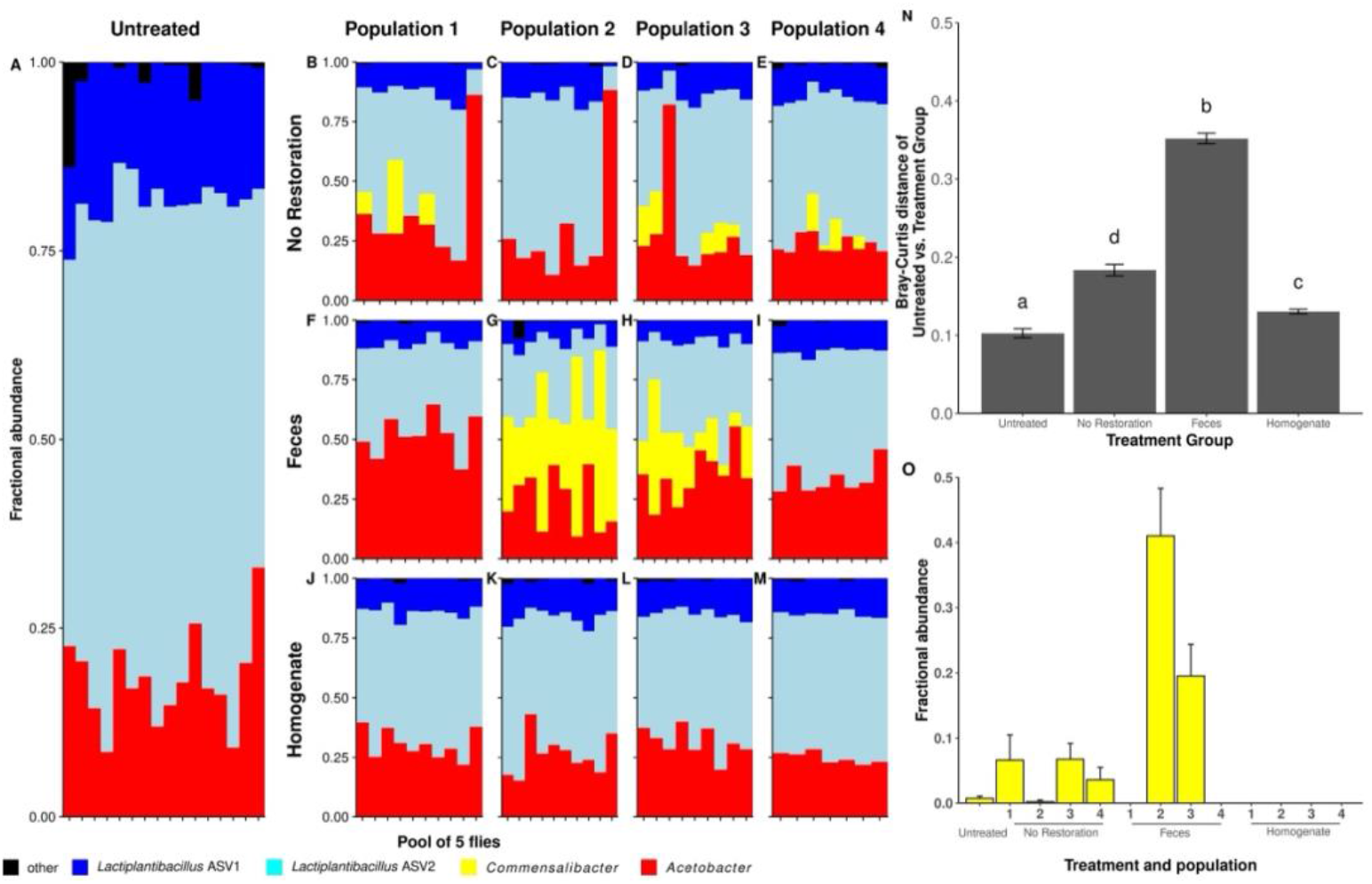
Microbiota composition after restoration in Wolbachia-colonized (W+) flies. Taxonomy plots of sequences clustered at the ASV-level for **A)** Untreated flies or **B-M)** flies from four replicate populations that each received restoration treatments. **N)** Bray-Curtis distances between the Untreated population and each restoration treatment (replicate populations are binned). Letters above bars identify statistically significant groups (p < 0.05). **O)** Relative abundance of a *Commensalibacter* ASV that was differentially abundant across treatments.

### Whole-body adult homogenates produce a native-like microbiota in flies following microbiota disruption

We also evaluated how different microbiota treatment strategies influenced microbiota composition in flies where the microbiota was disturbed by antibiotic treatment. A common way to eliminate the reproductive manipulator *Wolbachia* from flies is to expose flies to the antibiotic tetracycline over multiple generations. Therefore, we applied the Feces, Homogenate, or No Restoration treatments to four replicated lineages of W-flies two generations following a 3-generation exposure to tetracycline (the two-generation washout reduces possible impacts of the antibiotic itself). Unlike the conventionally-reared W+ flies, *Commensalibacter* were not appreciably abundant in these W-, microbiota-disturbed populations. However, as with the conventionally-reared W+ flies, the flies’ microbiota was of low diversity (**FIG 3A**), varied with restoration treatment (PERMANOVA: F_3, 113_ = 87.50, R^2^ = 0.67, p = 0.001), and resembled the Untreated flies most closely if they were treated with Homogenate (**FIG 3N, FIG S2D-E**). ANCOM revealed that two ASVs varied significantly with treatment: *Acetobacter*, which were depauperate (**FIG 3O**), and *Escherichia-Shigella* ASVs that were consistently enriched (**FIG 3P**) in the No Restoration flies relative to Untreated flies, respectively. Taken together, these findings show that, as in conventionally-reared flies, homogenates of microbes maintain generational similarity in microbiota composition better than fly feces or the absence of a microbial treatment.

**Figure 3.**
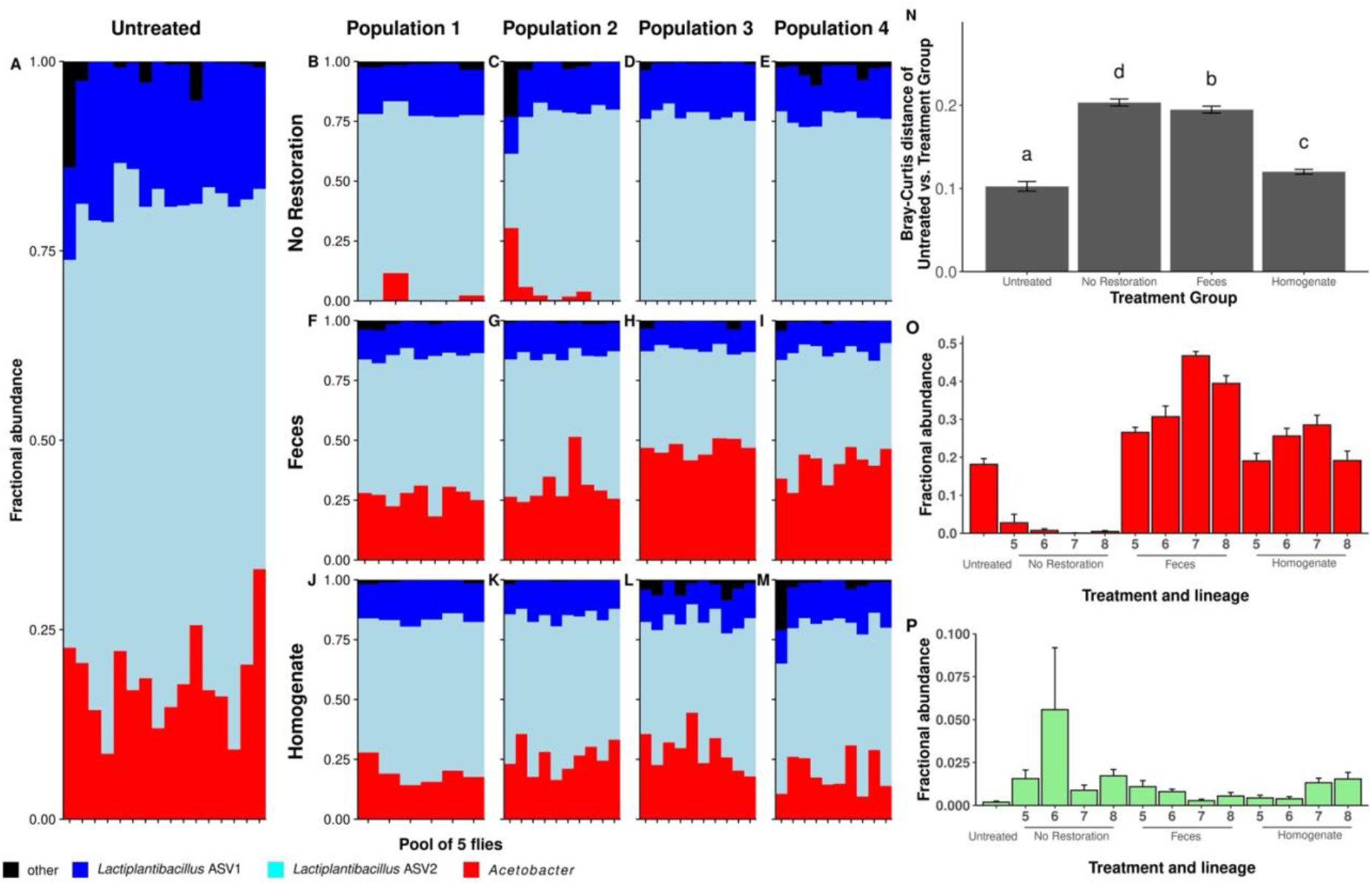
Microbiota composition after restoration in flies made Wolbachia-free by antibiotic treatment. Taxonomy plots of sequences clustered at the ASV-level for **A)** Untreated controls (age-matched W+ siblings that did not receive antibiotics) or **B-M)** antibiotic-treated flies from four replicate populations each that received one of three restoration treatments **N)** Bray-Curtis distances between Untreated flies and each restoration treatment (replicate populations are binned). Letters above bars identify statistically significant differences between treatments (p < 0.05). Relative abundance of **O)** *Acetobacter* and **P)** *Escherichia-Shigella* ASVs that were differentially abundant across treatments.

### Wolbachia alters the microbiota composition of flies

Finally, we tested whether the presence of Wolbachia influenced the microbiota composition of flies subjected to each treatment condition by comparing the replicated W+ and W-populations from each treatment group. There were significant differences in microbiota composition between W+ and W-flies in all treatments using all beta-diversity distance metrics (except for weighted Unifrac distances between No Restoration flies (PERMANOVA F _1, 67_ = 72.7, R^2^ = 0.52, p = 0.25), confirming that *Wolbachia* influences the microbiota composition of these flies (**TABLE 1, FIG S2G-I**). Overall, there was less variation in the microbiota composition of W+ and W-flies that received Homogenate treatment than from flies in other treatment conditions (**FIG 4A-F**).

**Figure 4.**
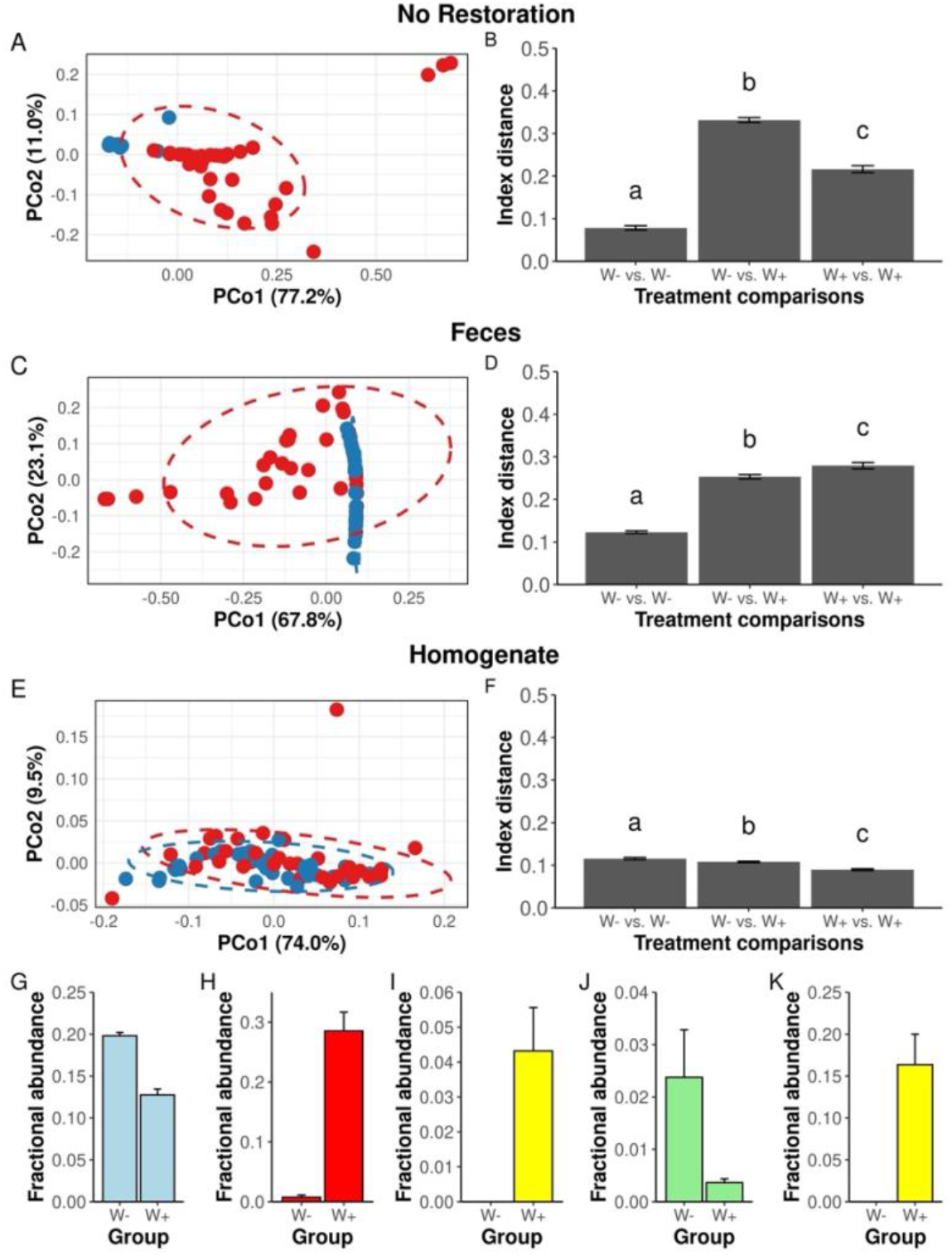
The effect of Wolbachia on microbiota composition in flies that received restoration treatments. **A**,**C**,**E)** Principal Coordinate Analysis (PCoA) ordinations of Bray-Curtis distances and corresponding **B**,**D**,**F)** beta-diversity distances of W+ (blue points) and W-(red points) flies reared under three microbiota restoration treatments. Letters above bars identify statistically significant differences in Bray-Curtis distances (p < 0.05). Mean relative abundances ± SEM of the following ASVs between W- and W+ flies in G-J) No Restoration and K) Feces treatments.

Homogenate-treated flies were also unique because *Wolbachia* reduced, rather than amplified, the beta-diversity distances between individual flies, and because no ASVs differed in abundance between Homogenate-treated W+ and W-flies. Conversely, a *Commensalibacter* ASV was more abundant only in *W+* flies that received No Restoration or Feces (**FIG 4G,K**), and *Wolbachia* influenced the abundance of ASVs assigned to *Lactiplantibacillus, Acetobacte*r, and *Escherichia-Shigella* only in flies that received No Restoration. Taken together, these findings are consistent with previous reports that *Wolbachia* can shape microbiota composition in flies and identify subtleties in these influences that may be tied to the microbial exposure of their hosts.

### Wolbachia influences fly locomotion

Having established a population of *Wolbachia-*free, microbiota-restored flies, we examined if *Wolbachia* contributed to fly activity, a trait which we have established expertise to measure. We compared climbing traits between the Untreated population and Homogenate-treated population 5, which had the most similar microbiota composition to the Untreated population (**FIG S3**). Flies colonized with *Wolbachia* climbed less distance and more slowly than W-flies (**FIG 5**). Conversely, W+ flies initiated more climbs than W-flies, which would be consistent with each climb occupying less time (**FIG 5**). These effects were specific to climbing because, consistent with previous comparisons of W+ and W-fly lines, *Wolbachia* did not influence overall fly movement (46). Taken together, these findings identify climbing functions as yet another *Drosophila* trait that can be influenced by *Wolbachia*.

**Figure 5.**
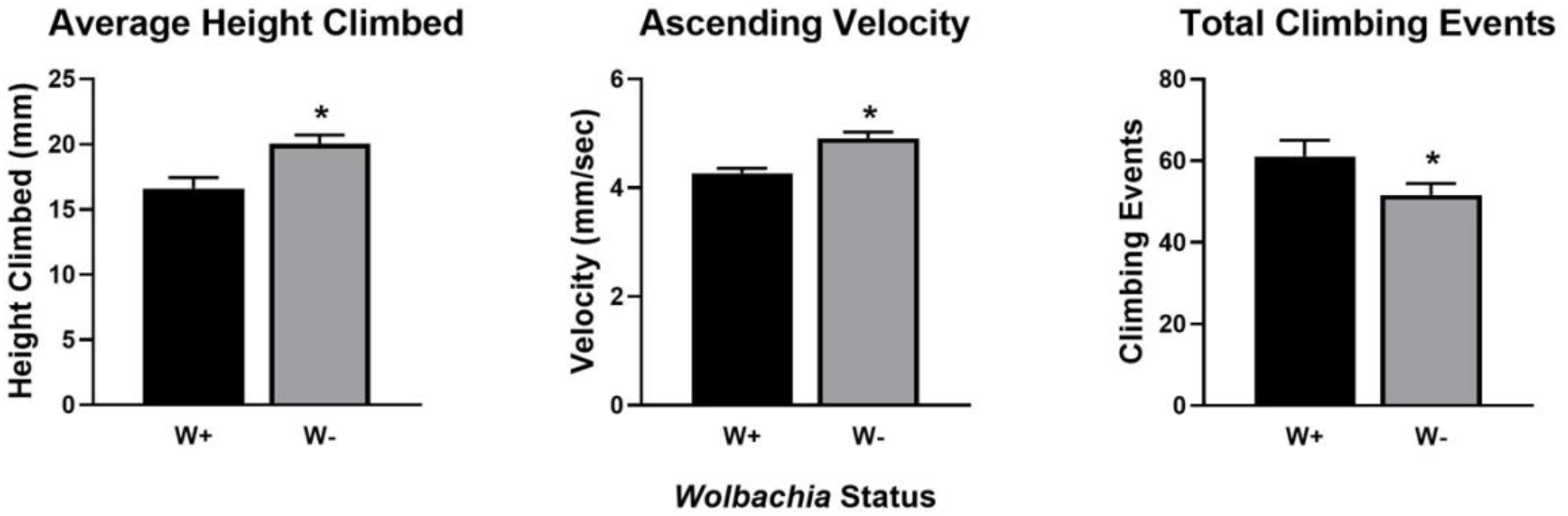
Wolbachia affects climbing ability and behavior. Bars represent mean ± SEM, * = p < 0.05.

## DISCUSSION

This study evaluated the efficacy of different microbiota restoration strategies in *Drosophila melanogaster* in the presence and absence of antibiotic-induced disruption of the gut microbiota. By comparing the microbiota composition between an untreated reference population and populations that were exposed to fly feces, whole-body homogenates, or no treatment diets, we found that flies fed a fly homogenate best resembled the untreated, “normal” condition flies of the preceding generation. These results enhance our understanding of microbiota resilience and show homogenate transfer can be a biologically relevant method for microbiota repopulation in experimental *Drosophila* models.

Consistent with previous studies in *Drosophila*, our antibiotic treatment significantly disrupted microbial community composition, reaffirming the impact of tetracycline on both *Wolbachia* clearance and broader microbiota alteration (47, 48). Flies in the No Restoration group exhibited the highest inter-individual variability, showing the limitations of relying on spontaneous environmental recolonization to recapitulate the microbiota of previous fly generations (8, 31). Feces transfer incompletely restored microbiota composition to that of previous generations relative to flies that received No Restoration. In contrast, flies fed diets inoculated with a fly Homogenate consistently exhibited microbial community structure that most closely clustered with the Untreated population of the three treatments we tested. These findings show that targeted microbial supplementation can enhance microbiota recovery and functional restoration (8, 49).

Previous findings help contextualize our conclusions regarding the differential outcomes of microbiota restoration methods in *Drosophila*. Inoculating antibiotic-treated flies is expected to normalize the microbiota, as the gut community is inconstant and sustained by dietary replenishment (8, 31). These characteristics likely explain why the microbiota of Homogenate-treated flies was more consistent across individuals compared to the No Restoration group.

However, it does not fully account for the unexpectedly high variability observed in the Feces group. One possible explanation is that this effect is specific to our study, which featured relatively low microbial diversity even for *Drosophila*. Studies with broader baseline diversity might yield different outcomes. Alternatively, feces may not serve as a uniform or reliable seeding source. Most viable bacterial cells in *Drosophila* reside in the foregut (50), and most do not survive gut transit (51). Moreover, the fecal microbiota often differs from the microbiota of dissected guts or whole flies, and may be dominated by taxa that are not typically abundant in the host, such as members of the Pasteurellales (52), Enterococci (53), or the “Enteric Bacteria Cluster” (54). As a result, fecal inoculation may introduce non-colonizing or transient microbes, making it more susceptible to priority effects or other ecological dynamics that hinder stable community establishment. Regardless of the underlying mechanism, our findings clearly demonstrate that inoculation with sibling homogenates more effectively recapitulates the Untreated control microbiota than fecal seeding.

*Wolbachia* presence emerged as another key factor influencing gut microbiota composition. Across all treatment groups, the microbial communities of W+ and W-flies differed, suggesting that *Wolbachia* may facilitate microbial colonization or promote coexistence of a specific set of commensals. We also observed that *Wolbachia* contributed to a specific set of fly locomotor phenotypes. These findings underscore the importance of accounting for *Wolbachia* status in experimental design and data interpretation, as it can serve as a biological confounder in microbiota studies. Although *Wolbachia*-mediated suppression of *Acetobacter* has been reported in multiple studies, this effect appears to be host genotype-dependent, complicating the generalization of findings—including our own. The relationship between *Wolbachia* and Acetic Acid Bacteria remains inconsistent across studies, with reports of positive (55-57), negative (47, 55, 58), and neutral (59) associations in *D. melanogaster* and other *Drosophila* species (60). Notably, causal interactions have been demonstrated in mosquitoes, where *Asaia*—a dominant Acetic Acid Bacterium—can suppress *Wolbachia* vertical transmission (61). These complex and context-dependent interactions highlight the need for further mechanistic studies to disentangle the ecological and evolutionary dynamics between *Wolbachia* and gut microbial communities.

Despite these insights, our study has several limitations. First, we focused exclusively on short-term microbiota restoration and did not assess the long-term persistence or functional consequences of the reestablished communities. Previous work has shown that microbiota in *Drosophila* can shift over time and require frequent environmental replenishment, raising questions about the stability of restored communities (8). Future research could incorporate longitudinal sampling, along with behavioral or physiological assays, to evaluate host-level outcomes over time. Second, while 16S rRNA sequencing enabled broad taxonomic profiling, it provides limited resolution for functional inference. Shotgun metagenomic sequencing would offer a more comprehensive view of microbial gene content, functional restoration, and interspecies interactions.

In conclusion, homogenate feeding was more effective in reestablishing the gut microbiota composition of an untreated *Drosophila* population than fly feces or no restoration treatment. These findings support the use of homogenate transfer as a standardized and biologically relevant strategy for microbiota restoration in fly research. They also provide empirical evidence that transplants composed of host-colonized microbes can yield different outcomes than those derived from waste products and transient organisms. Additionally, our results underscore the influential role of *Wolbachia* in shaping microbial communities and affecting behavior; highlighting the complexity of post-antibiotic microbiota recovery. Insights from this study may inform broader applications in microbiota-based interventions and experimental design across diverse animal models.

## DATA AVAILABILITY

Sequences from this study are available in the NCBI sequence read archive under accession XXXXX (to be released upon acceptance).

## ACKNOWLEDGEMENTS

Research reported in this publication was supported in part by the National Institute of General Medical Sciences of the National Institutes of Health under Award Number R15GM140388. The content is solely the responsibility of the authors and does not necessarily represent the official views of the National Institutes of Health.

## SUPPORTING INFO

**Figure S1.**
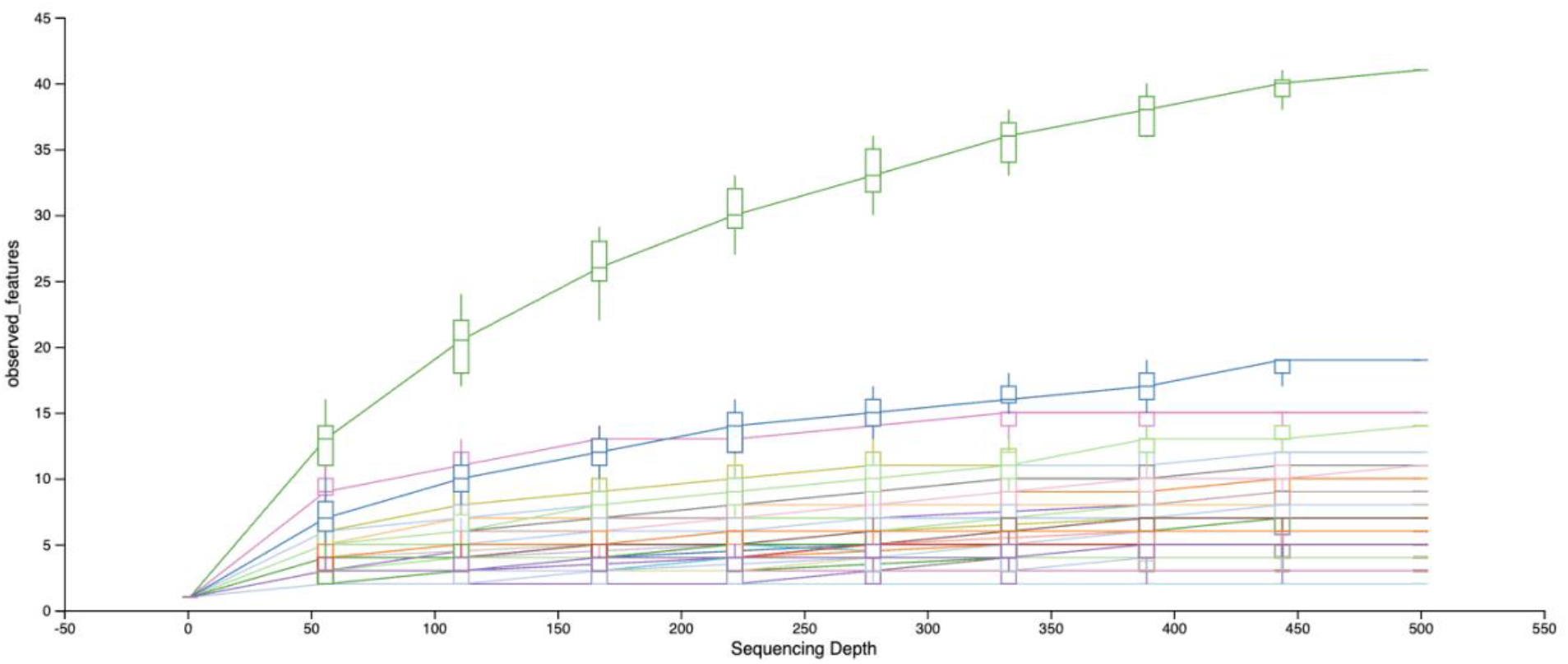
Rarefaction curve showing saturating sampling levels were approached. The curves show observed feature counts (y-axis) as a function of sequencing depth (x-axis). Most samples reached saturation by 500 sequences, supporting the chosen rarefaction depth for downstream diversity analysis.

**Figure S2.**
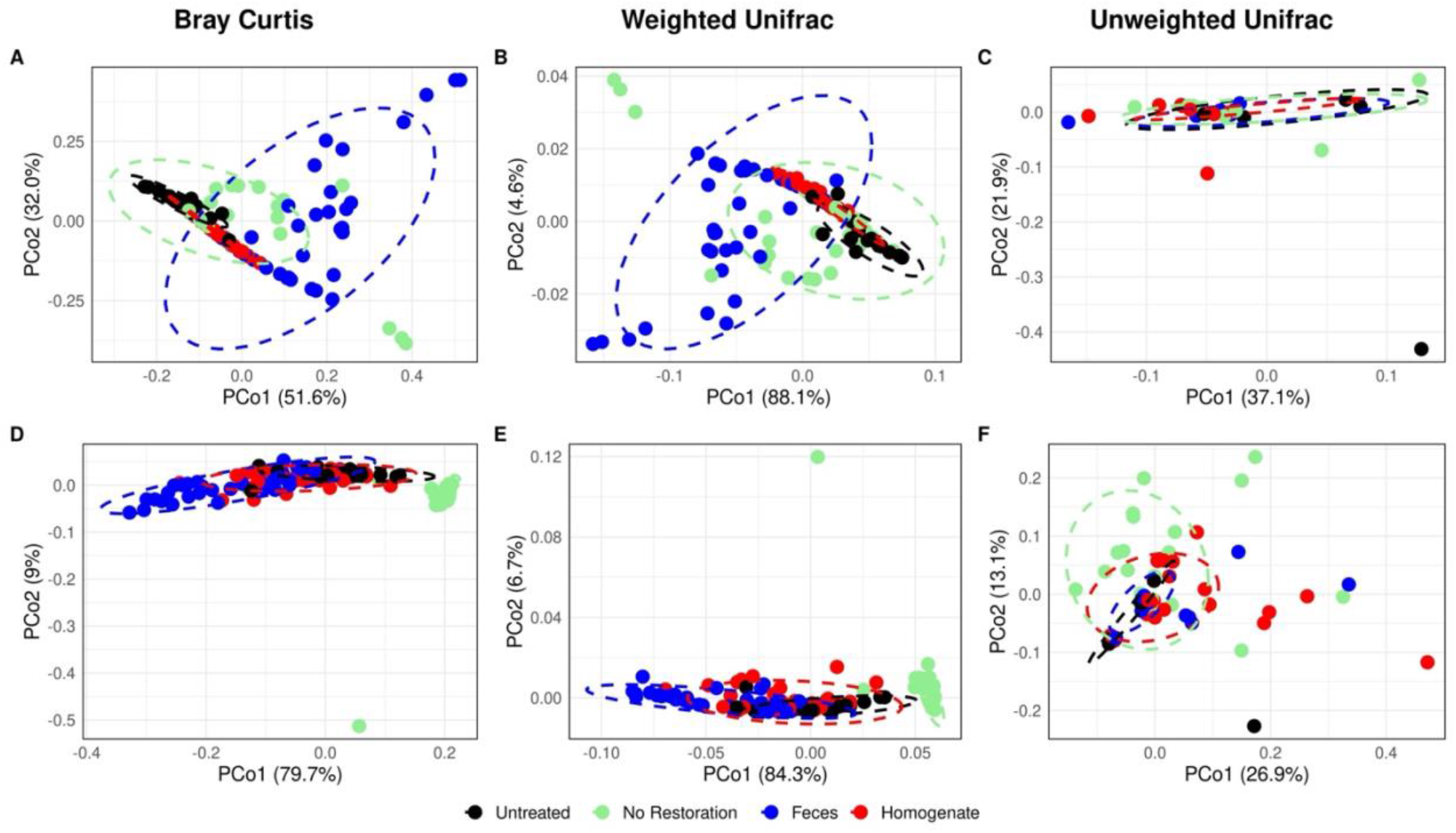
Principal coordinates analysis (PCoA) of W+ and W-flies. Principal Coordinates ordinations for A-C) W+ and Untreated, D-F) W- and Untreated flies.

**Figure S3.**
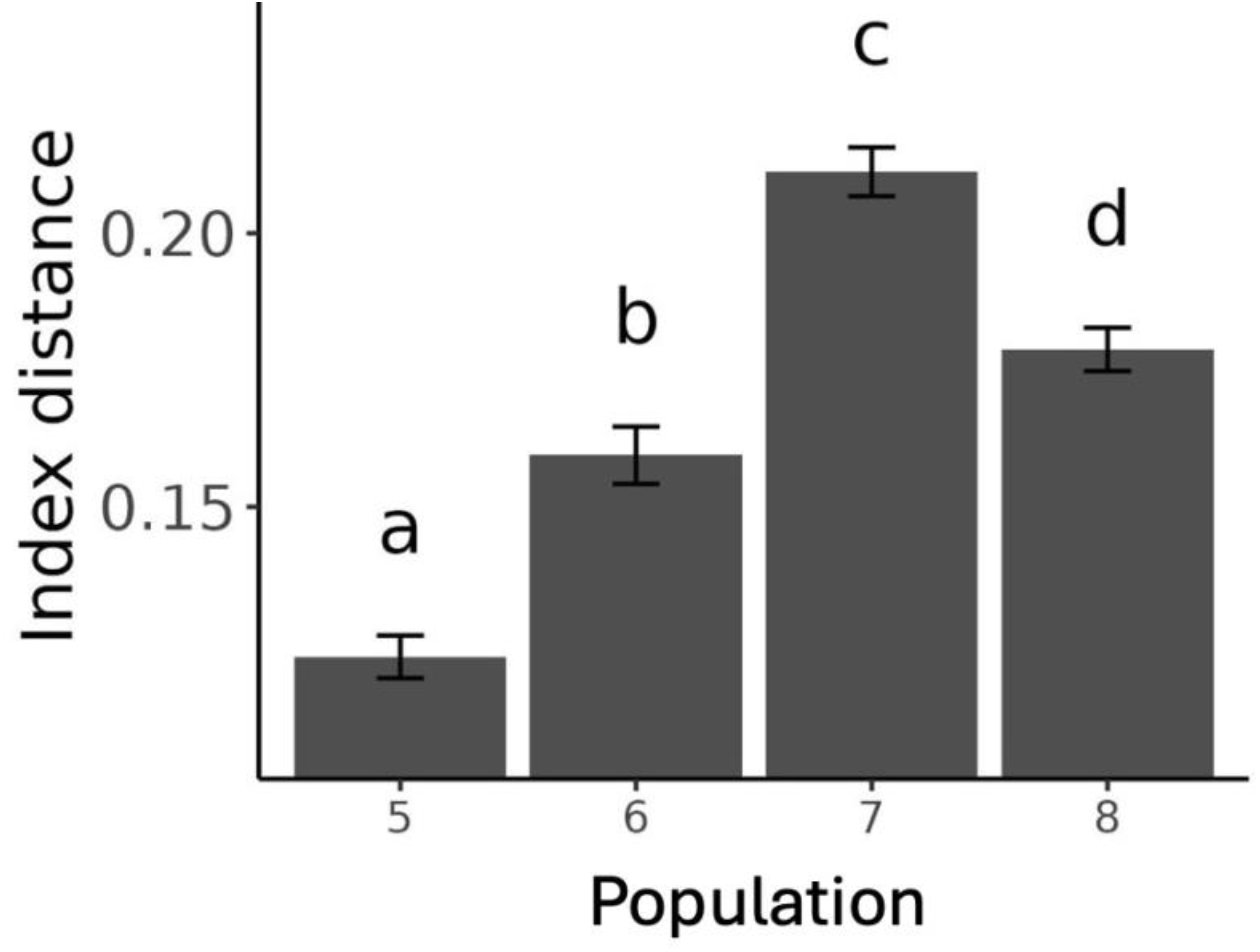
Bray-Curtis distances between W-Homogenate-treatment populations and Untreated flies. Compact letter displays show significant differences in the beta-diversity distances of W+ Populations 5-8 relative to Untreated flies. Population 5 was used in Figure 5 assays.

